# Cannabinoid CB_2_ receptor activation drives glucose uptake, shifting T cell metabolism

**DOI:** 10.64898/2026.07.07.736985

**Authors:** Robert S. Leddy, Hannah M. Phelan, Claire Connolly, Fabian Wehrmann, Desmond C. Winter, Lorraine Brennan, David O’Connell, Carol M. Aherne, Colm B. Collins

**Author notes:** **Correspondence**: Colm Collins Ph.D., Assistant Professor, UCD Conway Institute, University College Dublin, Belfield, Dublin 4, D04 V1W8, Ireland, E: Colm.

## Abstract

Cannabinoid receptor 2 (CB_2_R) is highly expressed on immune cells, but its role in T cell metabolism remains unclear. Here, we show that CB_2_R activation rapidly increases glucose uptake in human Jurkat T cells and drives a broader metabolic reprogramming away from glycolysis toward oxidative metabolism and the pentose phosphate pathway. Pharmacological CB_2_R activation increased mitochondrial mass, spare respiratory capacity, proton leak, and NADPH production, while CB_2_R inverse agonism produced the opposite effects. These metabolic changes were accompanied by upregulation of key pentose phosphate pathway enzymes, including GALT and TALDO1, and were abolished in CNR2-deficient cells, confirming receptor dependence. In primary human lamina propria mononuclear cells, CB_2_R signalling also influenced memory and gut-homing-associated T cell phenotypes, including integrin α4β7 expression. Together, these findings identify CB_2_R as a regulator of T cell bioenergetics and suggest that cannabinoid signalling may promote metabolic states linked to memory and tissue-homing functions in chronic intestinal inflammation.

**Brief Summary:**

- CB_2_R agonist-mediated increased glucose uptake in human T cells is lost in CNR2-deficient cells, suggesting that the response is receptor-dependent.
- CB_2_R activation rewires T-cell metabolism toward increased oxidative phosphorylation, spare respiratory capacity, mitochondrial mass, and pentose phosphate pathway metabolism, including higher NADPH and PPP-enzyme expression.
- In primary human intestinal immune cells, CB_2_R also promotes gut-homing and memory-associated T-cell phenotypes, including α4β7 and CD103 integrin induction ex vivo, suggesting a possible link to exacerbation of chronic intestinal inflammation.

## Introduction

T cell metabolism is a highly intricate and ever-changing process that is vital for the activation, differentiation, and maintenance of these immune cells. T cell activation is accompanied by extensive metabolic reprogramming that fuels clonal expansion and effector function. While naïve T cells rely primarily on catabolic metabolism including oxidative phosphorylation (OxPhos), antigen stimulation of the T cell receptor (TCR) drives a metabolic switch toward anabolic metabolism fuelled by aerobic glycolysis^1^. Even in the presence of adequate oxygen supply, this glycolysis by proliferating T cells is far less efficient than OxPhos at adenosine triphosphate (ATP) generation, in a process known as the Warburg effect^2^. Glycolysis supports rapid ATP generation and the high rate of T cell proliferation needed to mount an effective adaptive immune response. T cell activation increases glucose uptake 20-fold within 1 h^3, 4^, partly mediated by TCR-induced Glut1 translocation to the cell surface^5^.

In parallel, the pentose phosphate pathway (PPP) supplies ribose-5-phosphate for nucleotides and NADPH for reductive biosynthesis and antioxidant defence. Mitochondrial metabolism remains indispensable, however, as OxPhos and spare respiratory capacity (SRC) confer metabolic resilience under fluctuating nutrient and oxygen availability. Importantly, the balance between glycolysis and OxPhos also influences fate determination: effector T cells depend on glycolysis, whereas memory T cells maintain mitochondrial fitness, enhanced SRC, and oxidative metabolism to ensure long-term survival and recall capacity. Blockade of glycolysis also promotes Treg development and reduces the differentiation of CD4^+^ T cells into Th2^6^ and Th17^7^ phenotypes. Additionally, the knockout of Glut1 in T cells impairs glucose uptake, glycolysis, proliferation, and effector function in Th1, Th2, and Th17 cells, while Treg differentiation remains unaffected^8^. This is likely due to Tregs expressing low levels of Glut1 and being heavily dependent on OxPhos for their metabolic programming, so much so that inhibition of this metabolic pathway results in a significant impairment in the percentage of CD4^+^ T cells skewed towards the Treg lineage^9^.

Increasing evidence suggests that such pathways are not solely regulated by TCR and costimulatory signals, but also by extracellular modulators including the endocannabinoid system. Cannabinoid receptor 2 (CB_2_R), a Gi/o-coupled receptor highly expressed on immune cells, has been implicated in regulating cytokine production, cell migration, and inflammatory tone^10^. Mechanistically, CB_2_R engagement reduces intracellular cAMP, activates PI3K/Akt and ERK signalling, and induce AMPK activity. These pathways converge on metabolic checkpoints: PI3K/Akt promotes glycolytic flux and anabolic growth, ERK supports mitochondrial biogenesis and ROS regulation, and AMPK favours oxidative metabolism and spare respiratory capacity. Through these mechanisms, CB_2_R activation has the potential to remodel T cell metabolic programs, shifting the balance between glycolysis, OxPhos, and PPP activity. In this study, we investigate how CB_2_R signalling regulates core T cell metabolic pathways, with particular focus on glycolysis, OxPhos, PPP activity, and SRC. By delineating how CB_2_R engagement influences the balance between effector and memory programs, we aim to identify novel therapeutic opportunities to fine-tune T cell responses in health and disease.

## Methods

### Jurkat T Cell Culture

Jurkat T cells were obtained from the American Type Culture Collection (ATCC, Manassas, Virginia, USA) and maintained in RPMI-1640 medium containing L-glutamine (ThermoFisher Scientific; Cat No. 61870-010) supplemented with penicillin-streptomycin (ThermoFisher Scientific; Cat No. 15140122) and fetal bovine serum (FBS) (Merck; Cat No. F9665) to a final concentration of 1% and 10%, respectively. Cells were incubated at 37°C (95% O_2_/5% CO_2_) and passed every 2-3 days. *CNR2^-/-^* inhibitory CRISPR (CRISPRi) Jurkat T cells were cultured similarly, but with the added presence of 800μg/ml hygromycin (ThermoFisher Scientific; Cat No. 10687010) and 1μg/ml doxycycline hyclate (Tocris BioScience; Cat No. 4090).

### Fluorescent Glucose Analogue Uptake Assay

Jurkat T cells (1 × 10^6^ cells/ml) were seeded in pre-warmed supplemented RPMI-1640 in clear, sterile, flat-bottom 96-well plates and treated in triplicate with JWH-133 or GP-1a. Vehicle controls received glucose-free medium alone. Cells were incubated at 37°C for the indicated times, then pelleted (400 × g, 5 min, room temperature), resuspended in glucose-free medium, and labelled for 10 min with 100 µM 2-NBDG (Thermo Fisher Scientific, N13195). After transfer to white opaque, flat-bottom 96-well plates (Thermo Fisher Scientific, 437796), cells were washed with assay buffer/PBS, pelleted, resuspended in 100 µl assay buffer/PBS, and fluorescence was read immediately on a ClarioStar PLUS plate reader (BMG LabTech) at 485/535 nm.

### Seahorse XF Pro Analyser Assay

On day -1, 200μl of XF calibrant solution (Agilent Technologies; Cat No. 100840-000) was added to each well of the utility plate, before the XF Hydrobooster was placed on top of the utility plate and pushed downwards to tightly assemble. The sensor cartridge (Agilent Technologies; Cat No. 100840-100) was then lowered through the opening on the XF Hydrobooster into the utility plate, to be submerged in XF calibrant in order to be sufficiently hydrated. The plate was then placed in a non-CO_2_ incubator at 37°C and left overnight. On day_0_, 25μl of oligomycin, BAM15, and Rotenone/Antimycin A (Agilent Technologies; Cat No. 103772-100), reconstituted in 4ml assay medium, were supplied to injection ports A, B, and C of the sensor cartridge at a final concentration of 1.5μM, 2.5μmM and 0.5μM, respectively. The utility plate filled with XF calibrant and capped with the sensor cartridge was positioned in the Seahorse XF Pro (Agilent Technologies), and signal calibration was performed. In the meantime, Jurkat T cells were plated at a density of 1x10^5^ in 100μl of supplemented RPMI-1640 into a clear, sterile, flat-bottom 96-well plate. For the indicated times and in biological and technical triplicates, the cells were exposed to varying concentrations of the synthetic CB_2_R agonist JWH-133 or inverse agonist GP-1a freshly prepared from 10mM stock solutions in glucose-free RPMI-1640 media. A subset of cells were activated using a combination of anti-CD3 (BioScience; Cat No. 16-0037-85) and anti-CD28 (BioLegend; Cat No. 30293) human antibodies at a concentration of 5μg/ml. Vehicle-treated cells received supplemented media in the place of any cannabinoid. After treatment, the cells were transferred to a pre-warmed poly-d-lysine-coated, flat-bottom XFe96 microplate (Agilent Technologies; Cat No. 103792-100). 50μl of assay medium (Agilent Technologies; Cat No. 103576-100) supplemented with XF 1.0M glucose solution (Agilent Technologies; Cat No. 103577-100), XF 100mM pyruvate solution (Agilent Technologies; Cat No. 103578-100), and XF 200mM glutamine solution (Agilent Technologies; Cat No. 103579-100) were added to each of the four corner wells, before the plate was centrifuged gently at 200 x g for 5 minutes to allow the cells to attach to the bottom of the wells. The supernatant from each well was removed gently, and each well was made up to 175μl with assay medium. The culture plate was then incubated in a non-CO_2_ incubator at 37°C for 60 minutes prior to the assay. Once calibration completed, the culture plate was introduced and the acquisition programme was run. Data analysis was complete using the Wave Software (Agilent Technologies).

### NADPH Determination

In 100µl of pre-warmed supplemented RPMI-1640, Jurkat T cells were plated at approximately 2x10^6^ cells/ml into a clear, sterile, V-bottom 96-well plate. For the indicated times and in biological and technical triplicates, the cells were exposed to 1μM synthetic CB_2_R agonist JWH-133 or 10 μM inverse agonist GP-1a in order to assess the effect of CB_2_R signalling on the rate of NADPH production in Jurkat T cells. Vehicle-treated cells received supplemented media in the place of any cannabinoid treatment. Following treatment, the cells were pelleted by centrifugation, washed in ice-cold PBS and resuspended in 200μl of the provided NADP/NADPH lysis buffer. NADPH levels following CB_2_R manipulation was determined using the Amplite® Fluorometric NADPH Assay kit “Red Fluorescence” (AAT Bioquest; Cat no. 15262) according to the manufacturer’s instructions, to quantitatively analyse the NADPH content of lysates at 540/590nm with a ClarioStar PLUS plate reader (BMG LabTech) relative to the NADPH standard curve.

### Measurement of Enzymatic Concentration Levels

In 1ml of pre-warmed supplemented RPMI-1640, Jurkat T cells were plated at 2.5x10^6^ cells/ml into a clear 24-well plate. In biological and technical triplicates, the cells were exposed to 1µM synthetic CB_2_R agonist JWH-133 or 10µM CB_2_R inverse agonist. Vehicle-treated cells received supplemented media in the place of any cannabinoid treatment. Following the indicated treatment times, the cells were pelleted and the supernatants were collected for future applications. The cell pellet was washed three times in ice-cold PBS and lysed using 150μl ready-to-use RIPA Lysis Buffer (ThermoFisher Scientific; Cat No. 89900) supplemented with cOmplete™ EDTA-free Protease Inhibitor Cocktail (Roche; Cat No. COEDTAF-RO). A BCA Assay was used to quantify the protein concentration according to the manufacturer’s instructions, with 200μg of protein being used for each sample. Human galactose-1-phosphate uridylyltransferase (GALT) and human Transaldolase (TALDO1) activity were assessed via enzyme-linked immunosorbent assay (ELISA) according to the manufacturer’s instructions (AssayGenie; Cat No. HUDL01149 and HUFI00548, respectively). The signal was measured by determining the absorbance levels of each sample at 450nm using the SpectraMax M3 (Molecular Devices) microplate reader relative to the standard curves created.

### Measurement of Enzymatic Expression Levels

In 1ml of pre-warmed supplemented RPMI-1640 or glucose-free RPMI-1640 media, Jurkat T cells were plated at approximately 2.5x10^6^ cells/ml into a clear 24-well plate. For 24 h and in biological triplicates and technical duplicates, the cells were exposed to 1µM synthetic CB_2_R agonist JWH-133 or 10µM CB_2_R inverse agonist, freshly prepared from 10mM stock solutions in supplemented RPMI-1640 or glucose-free RPMI-1640 media, respectively. Vehicle-treated cells received the respective media in both cases, in the place of any cannabinoid treatment. RNA extraction and purification from the cells was carried out using the RNeasy Mini kit according to the manufacturer’s instructions, and converted to cDNA using a high-capacity cDNA Reverse Transcription Kit. Relative quantification of mRNA expression was performed using Taqman Gene Expression Assays and the QuantStudio 12K Flex Real-Time PCR System, in order to assess the expression levels of *GALT* (ThermoFisher Scientific; Hs00377666_m1) and *TALDO1* (ThermoFisher Scientific; Hs00997203_m1) +/- glucose. Relative mRNA expression of *UCP1* (ThermoFisher Scientific; Hs01084772_m1), *UCP2* (ThermoFisher Scientific; Hs01075227_m1), and *UCP3* (ThermoFisher Scientific; Hs01106052_m1) was measured by TaqMan qPCR on a QuantStudio 12K Flex system, with 18S rRNA as the endogenous control.

### Measurement of Mitochondrial Mass

Jurkat T cells were plated at approximately 1x10^6^ cells/ml in a 96-well plate in 100µl of pre-warmed supplemented media. Cells were treated in triplicate with varying concentrations of the synthetic CB_2_R agonist JWH-133 or inverse agonist GP-1a at 37°C for 24 h. Following incubation, the cells were centrifuged for 5 minutes at 1500rpm. Live cells were stained using Live/Dead Fixable Aqua dye in the dark for 30 minutes on ice. Cells were pelleted by centrifugation at 1500pm for 5 minutes and washed once in PBS. The cells were pelleted again and resuspended in 100μl of MitoTracker™ Green FM Dye (ThermoFisher Scientific; Cat No. M46750) dye, incubated at 37°C for 60 minutes protected from light. Cell fluorescence was immediately analysed using the Cytoflex LX.

### Metabolomic Analysis

Jurkat T cells were seeded in supplemented RPMI-1640 at 4x10^6^ cells/ml in 24-well plates and treated in technical quadruplicate with JWH-133 (1 µM), GP-1a (10 µM), or vehicle control for 48 h. Following incubation, cells were pelleted at 5000rpm, washed in ice-cold PBS, and subjected to targeted LC-MS/MS using the MxP® Quant500 assay (Biocrates Life Sciences, Innsbruck, Austria). Statistical analysis was performed in MetaboAnalyst 5.0 with Benjamini-Hochberg FDR correction, and group comparisons were made by one-way ANOVA with Tukey post hoc testing. Heatmaps and dot plots were prepared in MetaboAnalyst and GraphPad Prism 10, respectively.

### Human Lamina Propria Mononuclear Cells (LPMC)

Human intestinal tissue was collected from St. Vincent’s University Hospital (IRB Protocol No. RS19-068). Nine samples were obtained from both IBD patients and non-patients. The samples were cut longitudinally along the mesentery and all connective tissue and fat was removed. The luminal contents were washed in PBS, before the tissue was cut into short segments of approximately 12mm. These segments were placed in individual falcon tubes alongside 30ml isotonic solution (PBS supplemented with 400μl 0.5M UltraPure™ Ethylenediaminetetraacetic Acid (EDTA) (Invitrogen; Cat No. 15575-038) and 3ml 0.5M 4-(2-Hydroxyethyl)-1-piperazineethanesulfonic Acid (HEPES) (Gibco; Cat No. 15630-056)). The samples were vortexed at full speed for 5 minutes, until the solution had a transparent appearance. The tissue was then washed once in PBS, strained and blotted dry. In a petri dish, the tissue was minced with scissors and transferred to a 50ml falcon tube containing 20ml supplemented RPMI containing 0.75ml 0.5M HEPES and 25mg Collagenase I (Sigma; Cat No. SCR103). The tissue was incubated at 37°C for 60 minutes. Once complete, the tubes were quenched on ice and passed through a 70μm filter. The filtrate was then pelleted via centrifugation at 1,200rpm at room temperature and washed once in 5ml PBS. Cells were pelleted again and resuspended in 5ml PBS, before being counted and used as required.

LPMC were plated in supplemented RPMI-1640 (10^5^ cells/ml; 100µl/well) in flat-bottom 96-well plates and treated in triplicate with JWH-133 (1 µM), GP-1a (10µM), or vehicle control for 72 h at 37°C. For the final 4 h, cells were either activated with anti-CD3/CD28 antibodies (10 µg/ml) plus GolgiPlug™ or left unstimulated. Cells were then stained with the indicated surface markers and Live/Dead Fixable Aqua dye, with intracellular cytokine staining performed after fixation and permeabilization in activated samples. Samples were acquired on a CytoFLEX LX system.

### Statistics

Statistical analyses were performed using Student t test. Graphs are presented as means ± standard error of the mean (SEM) and were generated using GraphPad Prism 10 software. Values of P<0.05 were considered statistically significant.

## Results

T cell glucose metabolism is a pivotal aspect of T cell biology. Different T cell subpopulations utilise distinct metabolic pathways to meet their energy demands, with alterations in cellular metabolism having significant impacts on T cell fate determination. In addition to cellular immune responses, transcriptomic analysis identified glucose metabolism as being significantly regulated by CB_2_R signalling. To investigate this further, we assessed multiple aspects of T cell glucose metabolism in response to CB_2_R signalling manipulation, particularly focusing on the distinct metabolic pathways utilised by cannabinoid-treated cells, to test our hypothesis that CB_2_R signalling drives metabolic alterations in T cells that facilitates the development of long-lived, memory T cells.

### CB_2_R activation drives glucose uptake in human T cells

RNAseq data has previously identified a role for CB_2_R signalling in T cell glycolysis at a transcriptional level and as such we wanted to determine if CB_2_R activation altered glucose metabolism at a protein level^11^. We performed a fluorescent glucose analogue uptake assay in Jurkat T cells following 3 h, 6 h, and 24 h cannabinoid treatment. Uptake of the fluorescent glucose analogue 2-NBDG was quantified via fluorescence detection at Ex/Em of 485nm/535nm. At 3 h and 6 h, CB_2_R blockade and activation both significantly impacted on the rate of glucose uptake (**Figure 1A and B**), with a significant reduction with CB_2_R blockade and an increase relative to vehicle-treated cells with CB_2_R activation, respectively. This effect of CB_2_R signalling on glucose uptake was still observed at 24 h, albeit to a lesser extent and at a statistically insignificant level (**Figure 1C**). The effects of CB_2_R ligands on glucose uptake were abolished in CNR2^-/-^ Jurkat T cells, confirm the role of this receptor in mediating the alterations in the rate of glucose uptake observed (**Figure 1A**). To summarise, CB_2_R activation drives glucose uptake in T cells, in a process that is receptor-dependent and occurs rapidly.

**Figure 1:**
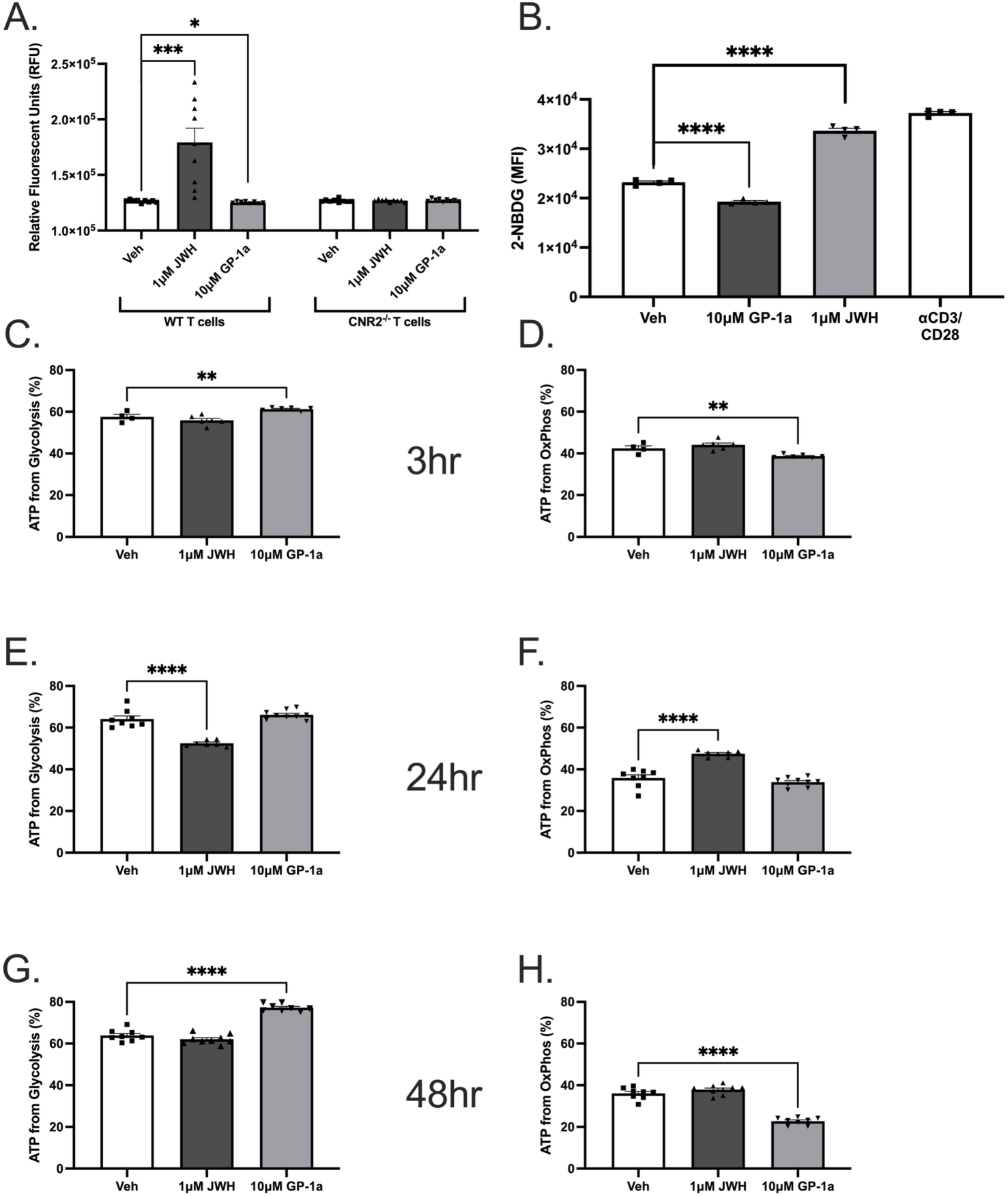
CB_2_R activation induces a rapid increase in glucose uptake in T cells, in a manner that is receptor-dependent. (A) Fluorescent 2-NBDG glucose analogue uptake assay in WT Jurkat T cells and CRISPRi CNR2^-/-^ Jurkat T cells following 3 h manipulation of CB_2_R signalling (N = 3, n = 3). (B) Fluorescent 2-NBDG glucose analogue uptake assay in WT Jurkat T cells following 6 h manipulation of CB_2_R signalling (N = 3, n = 3). (C) Fluorescent 2-NBDG glucose analogue uptake assay in WT Jurkat T cells following 24 h manipulation of CB_2_R signalling (N = 3, n = 3). Bar chart dots are representative of technical replicates. Statistical analysis performed using a two-tailed student’s T test, with values of significance being indicated as * p<0.05, ** p<0.01, *** p<0.001.

### CB_2_R signalling mediates a shift in the distinct metabolic pathways utilised by T cells during the process of glucose metabolism

Following its initial uptake, the manner in which glucose is utilised and managed within T cells has a major impact on regular cellular function, development, and differentiation. As a result, it was key for us to investigate the nature of glucose metabolism following CB_2_R manipulation, in order to get a further understanding of what is occurring within these cells following initial alterations in glucose uptake. To do this, we utilised Seahorse XF Analyser T cell metabolism assays to profile the glucose metabolism occurring in cannabinoid-treated Jurkat T cells at multiple timepoints. To measure the rate of glycolysis and OxPhos, ATP production was assessed from each pathway. CB_2_R blockade significantly increased the rate of glycolysis and decreased OxPhos across multiple timepoints **(Figure 2 A and C)**. In contrast, receptor activation significantly reduced glycolytic flux after 24 h cannabinoid treatment, with a corresponding increase in OxPhos **(Figure 2B)**. Taken together, these findings are suggestive of a major role for CB_2_R activation in the regulation of glucose metabolism in T cells, initially with regards to glucose uptake but also in mediating a metabolic shift away from glycolysis towards increased OxPhos. With increases in the rate of OxPhos being indicative of elevated mitochondrial function and SRC in T cells, we wanted to assess whether the metabolic alterations observed translated to such events. Both proton leak and SRC were assessed via the use of the Seahorse XF Analyser. Proton leak was significantly increased with CB_2_R activation and reduced with the inverse agonist GP-1a in treated Jurkat T cells **(Figure 2E)**. SRCs in treated Jurkat T cells were also significantly decreased at all timepoints tested following CB_2_R inhibition, whereas receptor activation led to an elevation in SRC which was only deemed significant after 24 h cannabinoid treatment **(Figure 2F)**. To assess what may be driving the alterations in proton leak observed within the mitochondria of cannabinoid-treated Jurkat T cells, RT-PCR was utilised to investigate any alterations in the mRNA expression of UCP 1-3, encoding for the UCP proteins known to regulate and induce proton leak^12^. Indeed, CB_2_R activation had a significant effect in increasing the expression of UCP2 and UCP3 relative to vehicle-treated cells **(Figure S1)**, whereas UCP1 remained unchanged **(Figure S1)**. Receptor blockade had no impact on the expression of any of the UCPs tested.

**Figure 2:**
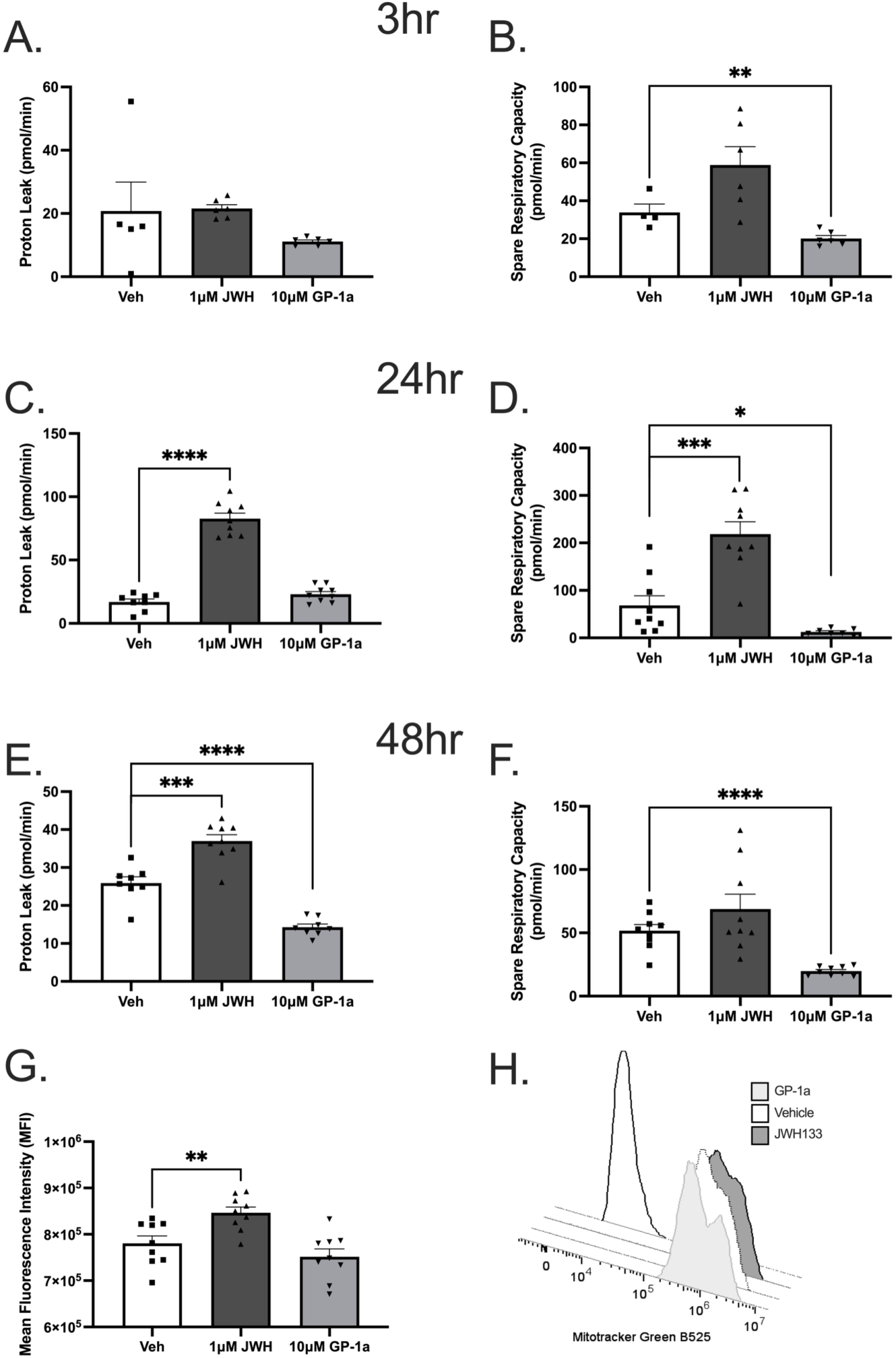
CB_2_R signalling regulates the distinct metabolic pathways utilised by T cells during glucose metabolism. Seahorse XF Analyser T cell metabolism assay assessing rates of glycolysis and OxPhos in Jurkat T cells following cannabinoid treatment for (A) 3 h (N = 3, n = 3), (B) 6 h (N = 3, n = 3, and (C) 48 h (N = 3, n = 3). Bar chart dots are representative of technical replicates. Statistical analysis performed using a two-tailed student’s T test, with values of significance being indicated as * p<0.05, ** p<0.01, *** p<0.001.

To assess whether the alterations in SRC following CB_2_R manipulation are due to increased mitochondrial biogenesis, we used flow cytometry analysis to observe any changes in mitochondrial mass following 24 h cannabinoid treatment, using the MitoTracker™ Green FM Dye. Activation of CB_2_R resulted in a significant increase in mitochondrial mass as indicated by MFI, relative to vehicle-treated cells **(Figure 2G, H)**. This is consistent with our previous Seahorse data assessing SRC in T cells following cannabinoid treatment. Inverse agonism had no significant impact on mitochondrial mass. Taken together, CB_2_R activation has a central role in the regulation of T cell glucose metabolism, by inducing the rapid uptake of glucose and mediating a shift towards increased mitochondrial respiration, characterised by increased OxPhos, enhanced SRC, and greater mitochondrial mass, all of which have been associated with the development of memory T cells.

### A metabolic reorganisation away from glycolysis towards the PPP occurs in T cells following CB_2_R activation

To further elucidate this metabolic shift occurring in T cells following CB_2_R activation, it was important to also investigate the expression levels of these key PPP-associated enzymes. We did so using RT-PCR to determine the mRNA expression levels of these enzymes in Jurkat T cells following 24 h cannabinoid treatment. In addition, due to the fact that CB_2_R signalling also drives the uptake of glucose into T cells, we measured GALT and TALDO1 mRNA expression in the absence of glucose using glucose-free culture media, in order to investigate the role of glucose in their expression. GALT expression was significantly increased in cannabinoid-treated cells following activation of CB_2_R relative to vehicle-treated cells, a result that was effectively abolished in the absence of glucose **(Figure 3A, C)**. An identical result was observed for TALDO1 **(Figure 3B, D)**. Receptor inhibition had no impact in altering the expression levels of these enzymes.

**Figure 3:**
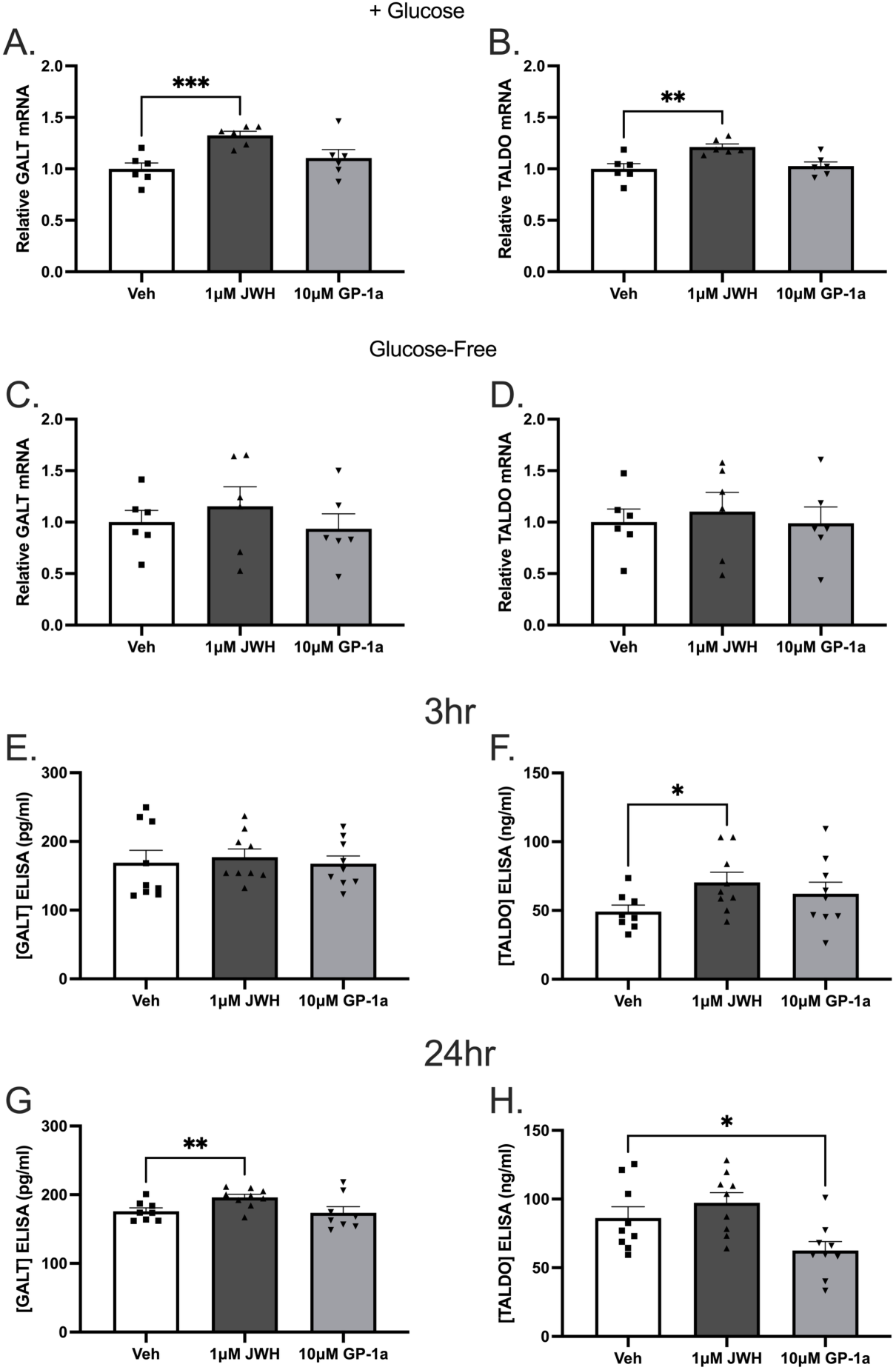
Both SRC and proton leak are altered in T cells following the manipulation of CB_2_R signalling. **(A)** Seahorse XF Analyser studies assessing the SRC of Jurkat T cells following 3 h, 24 h, and 48 h cannabinoid treatment (N = 3, n = 3). **(B)** Seahorse XF Analyser studies assessing the rate of proton leak in Jurkat T cells following 3 h, 24 h, and 48 h cannabinoid treatment (N = 3, n = 3). Bar chart dots are representative of technical replicates. **(E)** GALT concentration in Jurkat T cells following 3 h and (**G**) 24 h cannabinoid treatment investigated via ELISA assay (N = 3, n = 3)**. (F)** TALDO1 concentration in Jurkat T cells following 3 h and (**H**) 24 h cannabinoid treatment investigated via ELISA assay (N = 3, n = 3). Bar chart dots are representative of technical replicates. Statistical analysis performed using a two-tailed student’s T test, with values of significance being indicated as * p<0.05, ** p<0.01, *** p<0.001.

To further investigate a potential shift towards the PPP, we investigated the concentration of key PPP-associated enzymes in Jurkat T cells in response to acute and prolonged cannabinoid treatment via ELISA assay. The enzymes studied were GALT and TALDO1. GALT concentration was significantly increased following 24 h activation of CB_2_R signalling, whereas CB_2_R blockade had no impact across the two timepoints tested **(Figure 3E,G)**. On the other hand, TALDO1 concentration was found to be significantly elevated relative to vehicle-treated cells following acute CB_2_R activation for 3 h, a trend which was still apparent after 24 h but was deemed statistically insignificant. Prolonged receptor inhibition also led to a significant reduction in TALDO1 concentration, providing a contrast to the effects observed following receptor activation **(Figure 3F, H)**. The increased concentration of these key PPP-associated enzymes is consistent with the increases in NADPH observed, further suggesting a CB_2_R-mediated metabolic shift towards the PPP.

### Metabolomic analysis further reinforces the role for CB_2_R signalling in promoting a metabolic shift towards the PPP

Using LC-MS/MS, we were able to get a broad view of the alterations occurring within the T cell metabolome following receptor activation, across multiple metabolite classes. A total of 192 different metabolites from multiple compound classes, including acylcarnitines (N = 12), amine oxides (N = 1), amino acids (N = 17), amino acid-related compounds (N = 11), bile acid (N = 1), biogenic amines (N = 6), ceramides (N = 31), cholesterol esters (N = 1), diacylglycerol (N = 1), glycerophospholipids (N = 93), indoles derivative (N = 1), nucleobases-related compounds (N = 1), triacylglycerols (N = 15), and vitamins & cofactors (N = 1) were analysed. One-way ANOVA analysis (adj. p value ≤ 0.05) revealed that a total of 92 of these metabolites were significantly altered between treatment groups. The overall impact of CB_2_R activation is evident in the provided heatmap, with there being a distinctly metabolite profile being observed following receptor activation **(Figure 4)**. The metabolite classes most significantly altered were the amino acids, glycerophospholipids, and ceramides, with 65%, 59%, and 42% of each of these respective classes being significantly altered following manipulation of CB_2_R signalling **(Figure 5A,B)**. Analysis of these trends found that a number of PPP-associated amino acids were significantly increased following activation of CB_2_R signalling **(Figure 5C)**, which included histidine, phenylalanine, and tyrosine **(Figure 5D)**, consistent with our previous studies indicating a shift towards the PPP following receptor activation. With respect to the glycerophospholipids and ceramides, significantly altered members of these metabolite classes were plotted according to both acyl chain carbon length and their number of acyl chain double bonds, in order to assess whether either of these parameters could help determine the expression profiles of these metabolites. It was found that both the significantly altered glycerophospholipids **(Figure 5E)** and ceramides **(Figure 5F)** appeared to be generally decreased overall following CB_2_R activation relative to vehicle-treated cells. To summarise, LC-MS/MS metabolomic analysis allowed for a broad view of the metabolic alterations occurring in T cells following CB_2_R activation, with certain metabolite classes such as amino acids, glycerophospholipids, and ceramides being those most significantly altered. It is interesting to note that multiple PPP-associated amino acids are significantly increased following receptor activation, consistent with our previous studies which are suggestive of a metabolic shift towards the PPP following CB_2_R activation. Taken together, this data is strongly suggestive of a role for CB_2_R signalling in driving a metabolic shift T cell glucose metabolism, away from glycolysis and towards the PPP.

**Figure 4.**
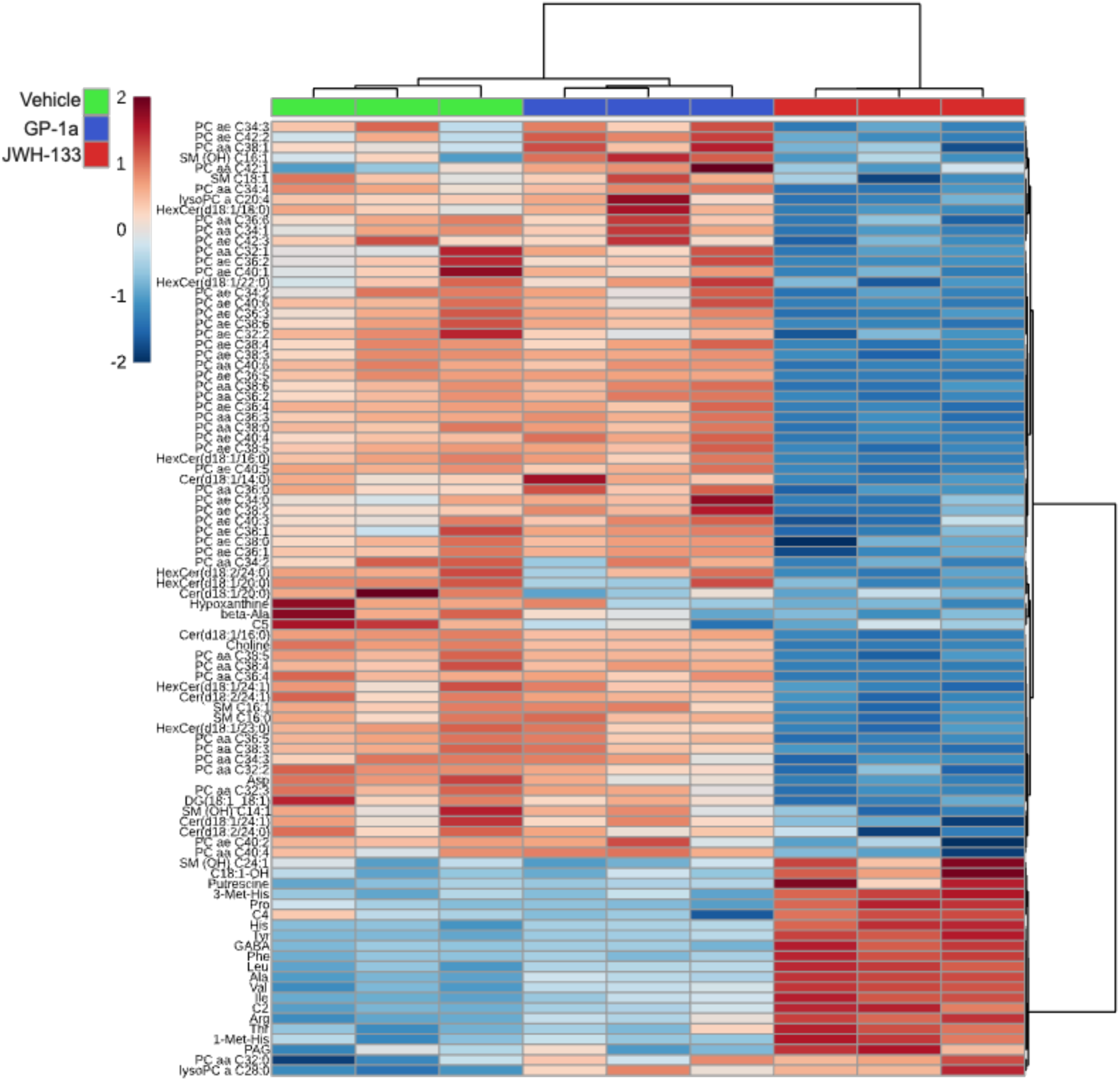
Changes in metabolic profiling in response to CB_2_R engagement. CB_2_R signalling is important in regulating the metabolomic profile of T cells, with receptor activation inducing distinct metabolic alterations. (A) Heatmap of the significantly altered metabolites (adj. p value ≤ 0.05) following 24 h CB_2_R inverse agonism or agonism (N = 92). The degree of positive or negative expression when comparing metabolites between treatment groups is expression is indicated by +2 (red) and -2 (blue).

**Figure 5:**
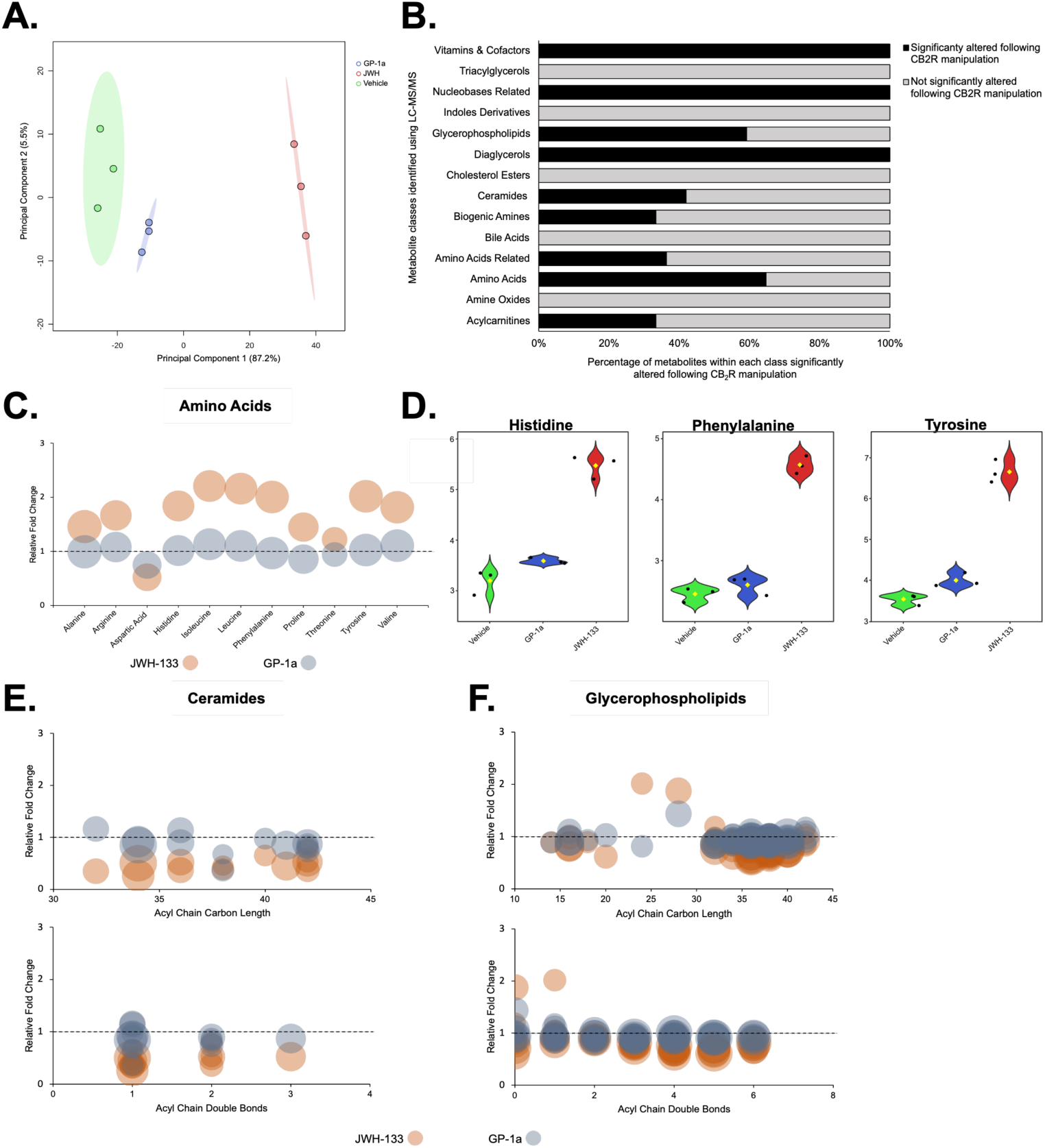
CB_2_R signalling alters T cell metabolome. (A) PCA score plot for the samples assessed via LC-MS/MS following cannabinoid treatment. R^2^ = 0.947 and Q^2^ = 0.866. (N = 3, n = 3). (B) Bar chart depicting the different metabolite classes altered in Jurkat T cells following 24 h CB_2_R manipulation, as assessed via Quant500 LC-MS/MS. (C) Significantly altered amino acids in Jurkat T cells following CB_2_R manipulation, plotted relative to vehicle-treated cells. Dot size is proportional to their degree of significance (-log(adj. p value)). (D) Significantly altered amino acids associated with the PPP. (E) Significantly altered ceramides in Jurkat T cells following CB_2_R manipulation, plotted relative to vehicle-treated cells. Dot size is proportional to their degree of significance (-log(adj. p value)). Ceramides are plotted according to acyl chain carbon length and acyl chain double bond number, respectively. (F) Significantly altered glycerophospholipids in Jurkat T cells following CB_2_R manipulation, plotted relative to vehicle-treated cells. Dot size is proportional to their degree of significance (-log(adj. p value)). Glycerophospholipids are plotted according to acyl chain carbon length and acyl chain double bond number, respectively.

### CB_2_R signalling may drive the development of human gut-homing memory T cells

To further assess whether CB_2_R signalling could potentially induce the development of memory T cells via metabolic reorganisation, a memory T cell conversion assay was conducted using isolated LMPCs from human intestinal tissue. Intestinal tissue from nine patients undergoing colectomy was collected, and LMPCs were isolated at a yield of approximately 1x10^7^ viable cells per collected sample which were frozen down prior to use. Using a logical gating strategy, we were able to filter out the CD3^+^ cells from the heterogeneous cell population isolated from the intestinal tissue, which accounted for approximately 86.7% of all live lymphocytes stained. From this cell population, CD4^+^ and CD8^+^ cell populations could be gated out, and as expected, the frequency of LPMC CD4^+^ was higher than LPMC CD8^+^ **(Figure 6A)**. We decided to focus mainly on CD4^+^ T cells due to their higher abundance within the isolated CD3^+^ population, and also their well-studied role in IBD, assessing the expression of particular immune cell markers on CD4^+^ T cells specifically. Our focus was on CD4^+^ memory T cells, using markers such as CD103, CD69, CD96, and CD45RO. Following receptor blockade, cumulative data found that expression of the marker CD103 was significantly decreased on isolated CD4^+^ T cells. This was also observed in concert with a trend towards a decrease in CD4^+^ CD103^+^ CD69^+^ T cells, a classical phenotype associated with tissue-resident memory T cells^13^. Conversely, activation of the receptor mediated no significant alterations in the generation of either CD4^+^ CD45RO^+^ CD95^+^ and CD4^+^ CD45RO^+^ CD69^+^ memory T cells **(Figure 6B)**, possibly due to the lack of sufficient biological replicates utilised in the assay. However, these immune cell markers are widely studied and utilised in the identification of memory T cells, particularly CD45RO.

**Figure 6:**
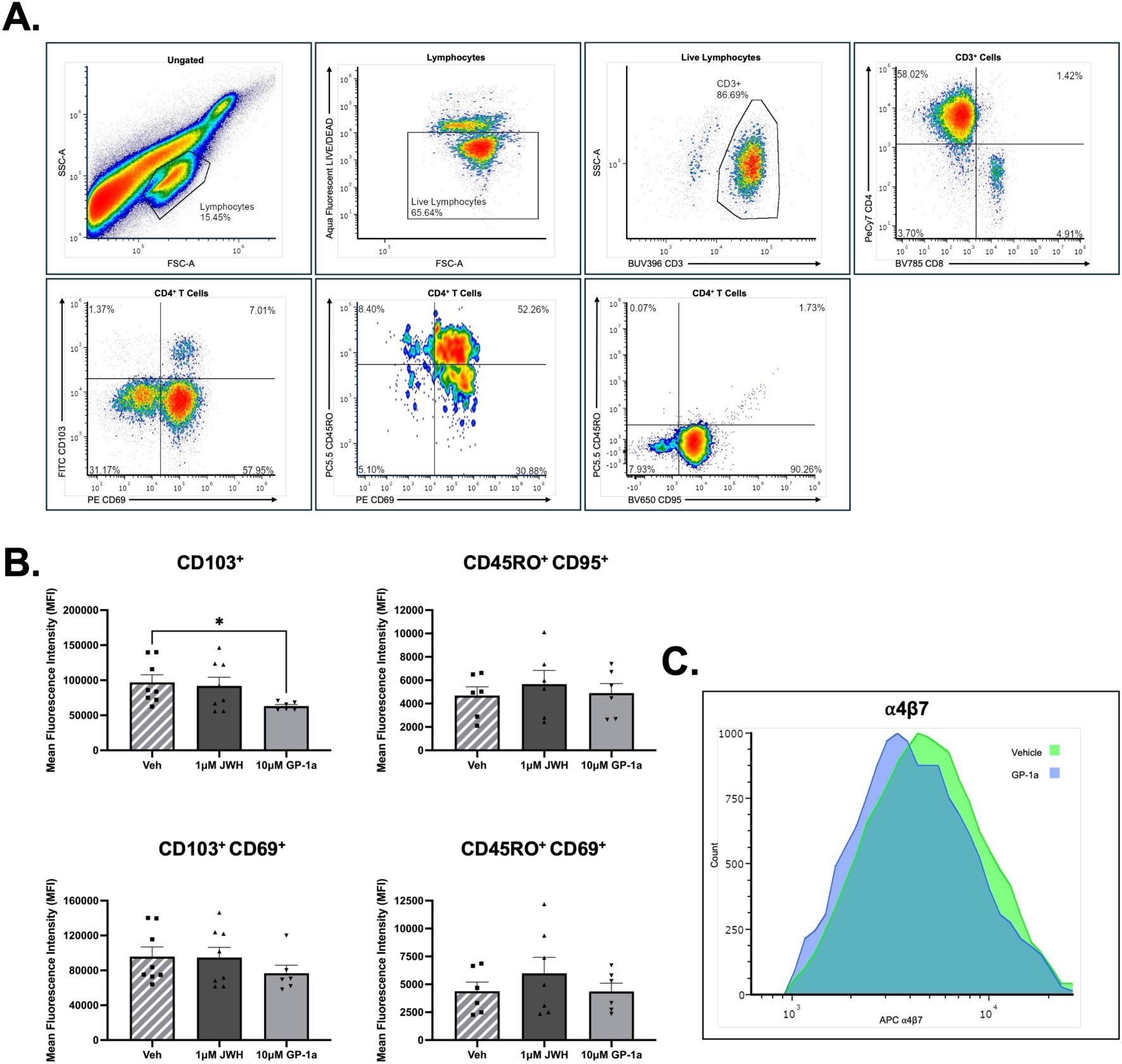
CB_2_R signalling alters the expression of memory T cell markers in human CD4^+^ T cells. **(A)** Flow cytometry gating strategy for the analysis of the expression of various immune cell markers on isolated human CD4^+^ T cells following 72 h cannabinoid treatment (N = 9, n = 3). Images provided are representative of merged FCS files. **(B)** Expression analysis of various immune cell markers, namely CD103, CD95, CD45RO, and CD69, on human CD4^+^ T cells following cannabinoid treatment assessed by means of MFI via flow cytometry. **(C)** Histogram overlay visually depicting the expression of integrin α4β7 on isolated human CD4^+^ T cells following CB_2_R blockade in comparison to vehicle-treated cells. The histogram provided is representative of the average MFI values of the merged files. Bar chart dots are representative of technical replicates. Statistical analysis performed using a two-tailed student’s T test, with values of significance being indicated as * p<0.05, ** p<0.01, *** p<0.001.

As previously reported, CB_2_R blockade significantly reduces the expression of the gut-homing integrin α4β7 on Jurkat T cells and their capacities in binding to MAdCAM-1. To assess whether the role of CB_2_R in this regard could be translated into human studies, we wanted to determine the expression of α4β7 on these isolated CD4^+^ T cells following cannabinoid treatment. Our results show that, consistent with our studies in Jurkat T cells, CB_2_R blockade decreased the expression of α4β7 on these human CD4^+^ T cells when compared to vehicle-treated cells **(Figure 6C)**. However, this decrease failed to reach statistical significance, again perhaps due to the significant degree of patient variability. In summary, our results indicate that CB_2_R signalling regulates the expression of integrin α4β7 on T cells and drives a metabolic reorganisation away from glycolysis and towards the PPP, potentially selecting for the development of gut-homing, memory T cells. Future studies are required to further consolidate whether CB_2_R signalling aids in the development of memory T cells.

## Discussion

Cannabinoid use is increasingly common among patients with inflammatory bowel disease (IBD), despite limited evidence that it improves underlying disease activity^14^ and evidence from a retrospective study suggesting potential harm^15^. We have previously demonstrated that cell-specific CB_2_R deletion or inverse agonism attenuates a model of human Crohn’s disease^11, 16^. This current study further informs how CB_2_R signalling may influence chronic intestinal inflammation. CB_2_R activation increased uptake of the fluorescent glucose analogue 2-NBDG in Jurkat T cells, whereas receptor blockade had the opposite effect. In the brain, CB_2_R activation stimulates glucose uptake in astrocytes and neurons. This is inhibited by CB_2_R-specific antagonists, consistent with a role for metabolic regulation^17^. Glucose metabolism is initially mediated by glucose uptake, primarily regulated by the transporter Glut1, the primary glucose transporter expressed in immune cells^5^. Glucose uptake via membrane-bound glucose transporters suppresses T cell proliferation, yet increased maturation and memory T cell formation^18^. Similarly, loss of *Glut1* suppresses effector T cell expansion and protected against piroxicam-induced colitis^19^ while blocking GLUT1 in a humanized UC model attenuated disease, in part by suppression of central memory CD4^+^ T cell development^20^. Single-sample gene-set enrichment analysis describes significantly higher rates of glycolysis in IBD patients’ intestinal tissues relative to controls^21^.

CB_2_R activation alters metabolism with increased intracellular glucose metabolised by the pentose phosphate pathway (PPP) at the expense of glycolysis. The apparent shift to PPP metabolism in CB_2_R^+^ T cells is consistent with previous studies of neuronal ischemia/reperfusion injury in which cannabinoids stimulated glucose metabolism through the PPP to maintain redox balance and energy conservation^22^. Memory T cells have lower glycolytic rates and higher levels of oxidative metabolism thereby limiting their growth but promoting their persistence^23^. In rheumatoid arthritis, a subset of glucose-6 phosphate (G6P) dehydrogenase G6PD^High^ cells divert glucose away from energy production toward the PPP, resulting in pathogenic CD4^+^ T cells with increased tissue invasiveness, accelerated conversion from naïve to memory phenotype and worsening inflammation^24^. Increased expression of TALDO1 alone would have minimal impact on nucleotide synthesis as the supply of G6P becomes rate limiting. GALT is a crucial enzyme in galactose metabolism, where it catalyses the reversible transformation galactose-1-phosphate into glucose-1-phosphate^25^. However, because CB_2_R activation also promotes glucose uptake and GALT upregulation to increase G6P synthesis from galactose, now G6P is no longer rate limiting. According to the principles of metabolic control, the pathway then becomes more highly sensitive to TALDO1 regulation, thereby allowing for a rapid increase in nucleotide synthesis without markedly decreasing concentration of metabolic intermediates in the main pathway (glutamine and aspartate) which would oppose the stimulation of macromolecule synthesis. PPP-metabolised glucose produces NADPH and metabolic intermediates required to generate macromolecules such as nucleotides, lipids and cholesterol for biosynthetic fitness^26–28^. NADPH is also important for maintaining a degree of redox balance within a cell, providing reduced glutathione and thioredoxins to neutralise ROS^29^. In lymphocytes, the NADP/NADPH ratio decreases from 1.4 in resting naïve cells to 0.2 in cells 4 days after stimulation, highlighting the requirement for the PPP and subsequent NADPH production in meeting the biosynthetic demands of clonal expansion^30^. We found CB_2_R signalling to significantly enhance the production of NADPH in cannabinoid-treated T cells, while also decreasing the NADP/NADPH ratios.

Consistent with the observed increases in NADPH following CB_2_R activation, receptor activation significantly enhanced TALDO1 concentrations. TALDO1 is a crucial enzyme in the PPP, facilitating the transfer of a dihydroxyacetone group from sedoheptulose 7-phosphate to glyceraldehyde-3-phosphate, forming erythrose 4-phosphate and fructose 6-phosphate^31^. This indicates a rapid shunt towards the PPP upon receptor activation. According to the principles of metabolic control laid out by Newsholme *et al.* (1985), if the flux through one branch is greatly in excess of the other, which in this case could be GALT in providing G6P, then the sensitivity of the low-flux pathway, namely PPP in this regard, becomes highly sensitive to pathway alterations such as TALDO1 regulation^32^. Pyruvate derived from the PPP is then preferentially shuttled into the TCA cycle and converted into acetyl CoA to fuel increased levels of OxPhos. *CNR2* CRISPRi cells confirmed the requirement for CB_2_R in mediating these effects, indicating the direct role for CB_2_R signalling in T cell glucose metabolism. These cells are more resistant to apoptosis, less metabolically active, and are smaller in size than effector cells, indicative of a cell that is less reliant on metabolism for clonal expansion^26^.

CB_2_R activation in T cells to induce a significant increase in mitochondrial mass. The increase in mitochondrial biogenesis and mass following receptor activation facilitates excessive OxPhos activity, mediating a metabolic shift away from glycolysis towards OxPhos. This shift is marked by enhanced SRC and increased mitochondrial efficiency, mediated by the expression of UCPs 2 and 3, driving proton leak and maintaining mitochondrial efficiencies through the potential reduction in mitochondrial ROS generation. As OxPhos and FAO are mitochondrial metabolic processes, alterations in mitochondrial mass allows memory T cells to effectively utilise these metabolic pathways to facilitate their long-term survival. These alterations in mitochondrial structure and function mediate distinct differences in the metabolic profile of a cell, and are distinct features distinguishing memory T cells from effector T cells. While effector T cells proliferate via mitochondrial fission, memory T cells undergo mitochondrial fusion, tightly assembling their mitochondria with the endoplasmic reticulum resulting in greater mitochondrial mass^33^. Memory T cells also exhibit higher SRC, the extra capacity of mitochondria to produce energy in response to increased stress or work, associated with prolonged cellular survival^33^. Memory T cells with their high SRC rates and greater mitochondrial mass more rapidly recall and respond to infections using this bioenergetic advantage^33^. In contrast, diminished mitochondrial mass of effector cells makes them heavily dependent on glycolysis due to their decreased ability to generate energy from alternative sources such as fatty acids via OxPhos.

Proton leak results in slow respiration, accounting for up to 25% of the basal metabolic rate^34, 35^. Proton leak reduces the electrochemical proton gradient, which operates in a cytoprotective manner in times of oxidative stress, in turn reducing superoxide production and lowering ROS levels^34, 36^. We found both SRC and proton leak mirror each other in cannabinoid-treated T cells, with receptor activation significantly enhancing both and CB_2_R inverse agonism reducing both. Generally, proton leak in mitochondria is induced by UCPs, particularly UCP2 and UCP3. These are proton channels expressed on the inner mitochondrial membrane which mediate proton leak to reduce ROS production. UCP2 is primarily expressed in immune cells, including T cells, with its expression tightly regulated and increasing significantly during T cell activation and proliferation, suggesting a role for this protein in the immune response^37, 38^. Both UCP2 and UCP3 are significantly increased in mRNA expression following CB_2_R activation, offering a potential mechanism through which CB_2_R signalling drives proton leak in T cells. In future, it would be useful to confirm that this increased proton leak and UCP upregulation translates to elevated mitochondrial ROS generation in T cells specifically following CB_2_R activation.

Using LC-MS/MS on cannabinoid-treated lysates provided unbiased screening of the T cell metabolome across multiple metabolite compound classes. We employed the MxP^®^ Quant500 system to measure concentrations of 500 metabolites across 26 different metabolite classes including glycerophospholipids, ceramides, and amino acids which were those most significantly altered following manipulation of CB_2_R signalling. The majority of the significantly altered glycerophospholipids were decreased upon CB_2_R activation, consistent with their role as substrates for the enhanced FAO and subsequent OxPhos observed in these T cells. In particular, we see a decrease in ceramides with CB_2_R activation, which have been previously linked with enhanced glucose utilisation and metabolism^39, 40^. Ceramide accumulation has also been linked to decreased mitochondrial respiratory chain activity and OxPhos, increased ROS generation, oxidative stress, and mitochondrial dysfunction. Ceramides directly inhibit mitochondrial complex III, to suppress mitochondrial OxPhos^41–43^.

To understand the changes in glycerophospholipid and ceramide profile in these T cells, both metabolite classes were plotted based on both their acyl chain carbon length and number of acyl chain double bonds. While most glycerophospholipids are decreased in T cells following CB_2_R activation, SM(OH) C24:1, with one double bond and an acyl chain carbon length of 24, was significantly increased. Its role in T cell biology is not well established and therefore warrants further study. CB_2_R activation increased PPP-associated amino acids, histidine, phenylalanine, and tyrosine. Histidine production is increased by catabolism of the PPP intermediate ribose 5-phosphate^44, 45^. Leucine treatment has also been associated with decreases in glycolysis, even in models where glycolysis substrates are sufficient^46^ and leucine-rich environments have been linked with a metabolic switch from glycolysis towards OxPhos^47^. These findings are consistent with the overall hypothesis that CB_2_R activation skews T cell metabolism from glycolysis to PPP and OxPhos to facilitate long-term survival. This shift would support memory T cell expression but this has not been proven unequivocally to date.

Human intestinal tissue-resident memory T cells express a range of markers such as CD45RO, CD69, CD95 and CD103^13^. CD103 or integrin αE forms a heterodimeric pair with integrin β7 which we have previously demonstrated is upregulated by CB_2_R activation^11^. CD103 binding to epithelial-expressed E-cadherin promoting the mucosal retention, provides long-lasting, localised immune surveillance^48^. We found a significant decrease in CD103^+^ CD4^+^ T cells, as well as a trend towards a decrease in CD103^+^ CD69^+^ CD4^+^ T cells with CB_2_R inverse agonism. These results suggest that CB_2_R inverse agonism may not only reduce the number of T cells trafficking to the gut, but also specifically targeting those that could become intestinal-associated tissue-resident memory T cells. This represent a novel therapeutic target for the management and treatment of IBD, given that proinflammatory tissue-resident memory T cells, marked by CD69 and CD103, are elevated in the intestinal mucosa of IBD patients, contributing to the chronic inflammation observed in IBD^49–51^. CB_2_R activation also increased the levels of CD45RO^+^ CD69^+^ and CD45RO^+^ CD95^+^ CD4^+^ T cells *ex vivo*. Increased levels of these cells is associated with disease flares and a more complicated disease course in patients^52, 53^.

Several limitations of this study must be acknowledged. Much of the metabolic work was performed in Jurkat T cells, and although these provide a tractable system for mechanistic study, they do not fully reflect the heterogeneity of primary human intestinal T cells. In addition, the link between the observed metabolic programme and bona fide memory-cell differentiation remains partly inferential. Future work will validate the mitochondrial and PPP findings at the protein and flux levels, directly measure ROS, and determine whether CB_2_R signalling promotes durable memory formation in primary T-cell systems *ex vivo* and *in vivo*.

Overall, these data identify CB_2_R as a regulator of T-cell metabolic organisation, linking cannabinoid signalling to glucose uptake, mitochondrial function, PPP activity and memory-associated phenotypes. These findings suggest that while CB_2_R activation may suppress effector T cell proliferation in the short term, it does so by supporting the shift to long-lived, gut-associated T-cell states that could exacerbate chronic intestinal inflammation. This provides a mechanistic framework for understanding why cannabinoid use may fail to improve IBD despite symptomatic benefit, and supports further investigation of CB_2_R blockade as a potential novel strategy to limit pathogenic T-cell persistence in chronic intestinal disease.

## Author Contribution

CBC conceived of the study. RL, and HP conducted experiments, generated reagents, analysed and interpreted data generated. CC and LB oversaw the metabolomic analysis and data interpretation. CA, FW, DCW and DOC guided the study design, provided reagents and helped with implementation. All authors contributed to refinement of the study protocol and approved the final manuscript.

## Abbreviations

(CB_2_R): cannabinoid receptor two
(CNR2): gene encoding cannabinoid receptor 2
(SRC): spare respiratory capacity
(FA): fatty acid oxidation
(G6P): glucose-6-phosphate
(GALT): galactose-1-phosphate uridylyltransferase
(IBD): inflammatory bowel disease
(LPMC): lamina propria mononuclear cells
(NADPH): nicotinamide adenine dinucleotide phosphate
(OxPhos): oxidative phosphorylation
(PPP): pentose phosphate pathway
(RA): retinoic acid
(ROS): reactive oxygen species
(TALDO1): transaldolase 1
(UCP1/2/3): uncoupling proteins 1, 2, and 3

